# The leaf cell arrest front is established by spatio-temporal control of cell sizes at division

**DOI:** 10.64898/2026.04.20.718680

**Authors:** Robert Kelly-Bellow, Rachel Prior, Nicola Trozzi, Mateusz Majda, Ross Carter, Matthew Hartley, Verônica A. Grieneisen, Athanasius F. M. Maree, Richard S. Smith, Michael W. Bevan

**Author notes:** joint first authors (Robert Kelly-Bellow, Rachel Prior, Nicola Trozzi). Senior authors (Richard S. Smith, Michael W. Bevan).

## Abstract

The sizes and shapes of organs are established by the combined actions of cell proliferation and cell growth. In plants, development of the determinate planar leaf is initiated by primordia formation and establishment of abaxial/adaxial polarity [1,2,3]. Lamina outgrowth is driven by cell division and growth along proximo-distal (PD) and medio-lateral (ML) axes [4], established by mutually repressive PD gradients of miRNA and target transcription factors [5,6,7,8]. These gradients generate proximal regions of competence for cell division and increased growth, with distal regions of reduced growth, endoreduplication and differentiation. The transition from proliferation to growth and differentiation is marked by a cell cycle arrest front, which moves basipetally during leaf growth, progressively restricting proximal proliferative zones as the leaf grows [9,10,11]. Intersection of proximal proliferation-promoting gradients with distal differentiation-promoting gradients may delineate the arrest front, but its dynamics remain poorly understood. We reasoned that mutants affecting cell proliferation patterns may provide insights into the formation, maintenance and dissolution of the arrest front. Spatio-temporal modelling of live imaging data of loss of function mutants of the regulatory peptidase DA1 and its E3 ligase activator Big Brother (BB), which increase cell proliferation [12,13], showed that these proteins effectively establish a threshold cell size at division as a function of distance from the base of the growing leaf and the duration of growth. Loss of BB and DA1 activities increased the persistence of cell divisions and dissolved the arrest front. This suggested that the arrest front emerges from the interactions of threshold areas of cell division with the cessation of division over time, and not from an independently-specified boundary.

## Results and Discussion

Leaf growth has been characterised at the cellular level using live imaging in growth chambers [14–16] coupled with MorphoGraphX [16–18]. This approach enables continued tracking of arrest front dynamics over an extended period of leaf growth. *DA1* family loss of function mutants have meristems with more cells and organs that continue to grow for a longer period than wt, due to a prolonged duration of mitotic cell divisions ([12,19–24]. The E3 ligase BB [12,13,19–21]) activates DA1 by monoubiquitylation, and loss of function of *BB* (the *eod1-2* T-DNA allele of *BB*) coupled with *da1-1* exhibit synergistic increases in leaf, petal and seed size [12]. The *da1-1eod1-2* double mutant was crossed with a *speechless* (*spch*) mutant expressing the pATML1::mCitrine-RCI2A epidermal-specific cell membrane marker to track epidermal cell outlines in the absence of stomatal proliferation [25]. *spch* has been shown not to influence leaf growth compared to wt in growth chambers and in petri dishes [15]. Figure 1A shows the *da1-1eod1-2* double mutant is comparable in size to the *da1-1eod-2 spch* triple mutant *pATML1::mCitrine-RCI2A* line. Continuous time-courses of images of leaf 1 from *da1-1 eod1-2 spch* and *spch* Arabidopsis seedlings grown in live imaging chambers [26] were captured using the methods of [15]. Leaf 1 was imaged every 6 hrs for the first 48 hrs, then every 8 hrs for the next 48 hrs, every 12 hrs for the next 48 hrs, and every 24 hrs for the final 48 hrs in three replicates for up to 300 hrs (12.5 days). Adaxial surfaces were segmented and cell location, shape and division patterns were quantified using MorphoGraphX [17,18]. Figure 1B (*spch*) and Figure 1C (*da1-1eod1-2 spch) s*how examples of cell segmentation in the growing leaves from 5-12 days after initiation (shortened to days). The *da1-1 eod1-2 spch* mutant had a larger leaf area and more cells compared to *spch* (Figure 1D). The range of areas of segmented cells was comparable between *spch* and *da1-1eod-2spch* at all time points in our dataset (Figure 1E), showing that the larger segmented area of *da1-1eod-2 spch* was due to increased cell proliferation. Figure 1F (*spch*) and Figure 1G (*da1-1eod-2 spch) s*hows the proportion of leaf cells with the highest areal growth rates in *spch* were at days 7 and 8, while in *spch da1-1 eod1-2* maximal growth rates were at day 10. Areal growth persisted in both lines until the last sampled time at day 12.

**Figure 1.**
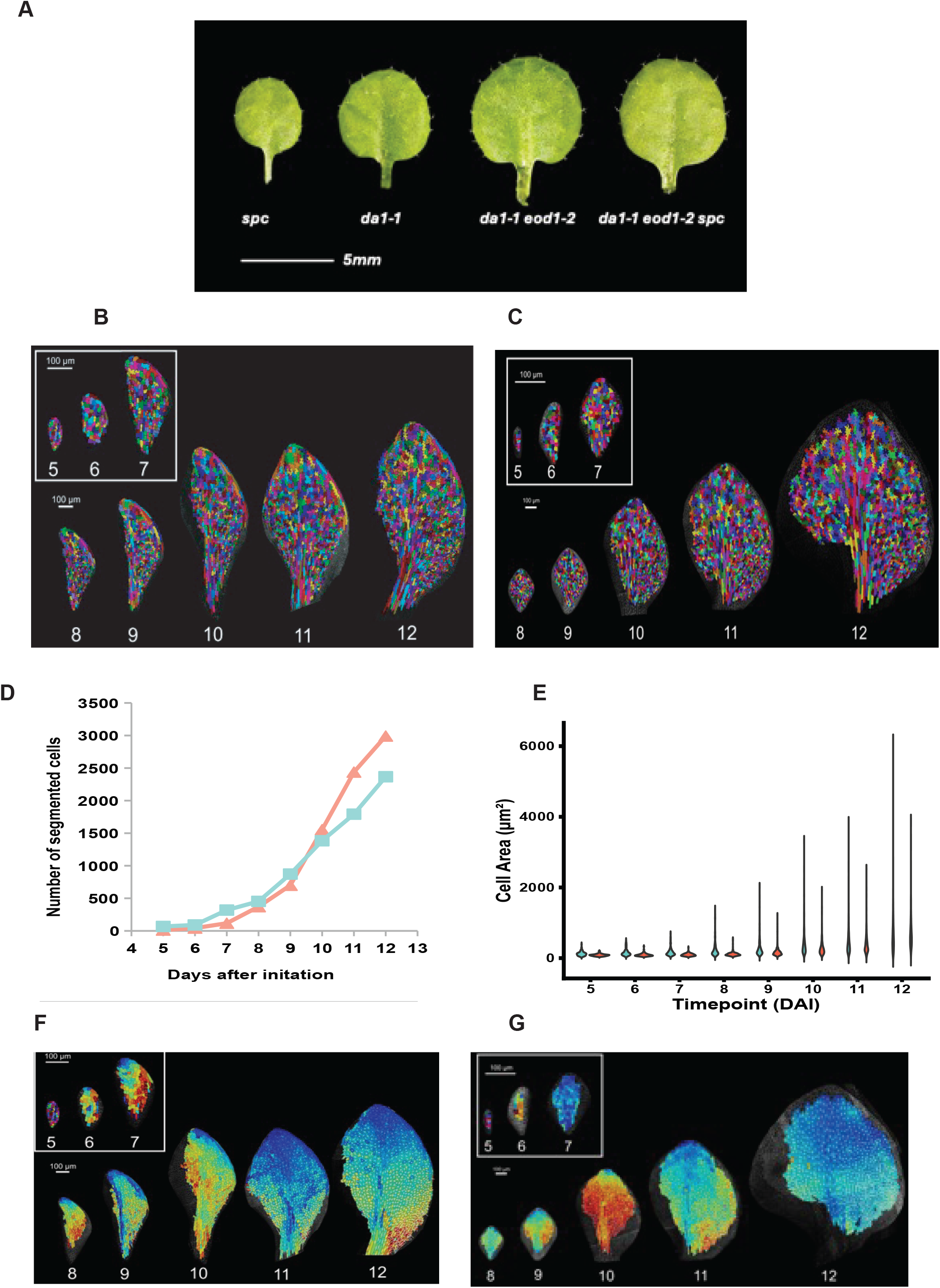
Leaf Growth, Cell Areas and Segmentation. A. Leaf 1 of plate-grown seedlings imaged at 14 days after germination. The typical larger leaf area of the double *da1-1 eod1-2* mutant is not altered by the *spch* mutant defective in stomata formation. B. Segmented *spch* leaf 1 at days 5-12 after initiation. The insert shows the different scaling used for smaller day 5,6,7 leaves. The scale bar is 100 μm. C. Segmented *da1-1 eod1-2 spch* leaf 1 at days 5-12 after initiation. The insert shows the different scaling used for smaller day 5,6,7 leaves. The scale bar is 100 μm. D. The graph shows numbers of segmented cells in *spch* (blue line) and *da1-1 eod1-2 spch* (red line). Note the increased cell numbers in *da1-1 eod1-2 spch* after day 10. E. The graph shows segmented cell areas during leaf growth. There was no significant difference between segmented cell sizes during growth of *spch* (blue) and *da1-1 eod1-2 spch* (red). F. Areal growth rates of *spch* leaves from days 5-12. G. Areal growth rates of *da1-1 eod1-2 spch* leaves from days 5-12. The scale bar shows high growth rates as red and low growth rates as blue.

### Cell proliferation in *da1-1 bb spch* occurs in larger cells and is more persistent

To understand the dynamics of proliferating cells in the mutant lines, we performed cell lineage tracking and proliferation analyses. We first took the number of cells that had divided between two time points as a percentage of all segmented cells at the later time point. Patterns of cell division are shown for *spch (*Figure 2A) and *da1-1eod1-2 spch* (Figure 2B). Between 7-8 days divided cells became limited to the proximal regions of leaf in both sets of mutants. In *spch* cell divisions became less frequent and ceased by day 12. This timing was consistent with a cell proliferation burst that becomes localised to proximal regions, defining a clear arrest front boundary, for example seen at day 8 in *spch* [8]. Cell division in *da1-1eod1-2 spch* at days 7-8 remained mainly in proximal regions as in *spch*, but after day 9 it became more widely distributed across the growing lamina compared to *spch*, with divisions continuing to occur all over the lamina until the final timepoint at day 12. In *spch* the numbers of proliferating cells decreased sharply after 8 days until no proliferating cells were found after 11 days. (Figure 2C). In *da1-1eod1-2 spch* cell proliferation decreased more slowly and many cells were still dividing at 12 days (Figure 2C). Overall, in 12 days of growth 370 cell divisions were detected in *spch* and 495 detected in *da1-1eod1-2 spch*, showing that together *DA1* and *BB* limit the numbers of cell divisions during leaf growth. To estimate the area at cell division, we used the average areas of parents and sum of daughter cell areas. For *spch* the average cell area at division was approximately 230 μm^2^ at all time points (Figure 2D). For *da1-1eod1-2 spch*, average cell area at division was comparable to *spch* from 6 to 8 days (*p* > 0.05), but between 9 to 11 days cell areas at division increased significantly in *da1-1eod1-2 spch* compared to *spch* (p < 0.001), with a very broadly distributed range of cell sizes at division, averaging approximately 600 μm^2^ at 12 days (Figure 2D). In *spch*, cells before division were smaller and usually elongated (Figure 2E). In *da1-1eod1-2 spch*, a typical dividing cell after 10 days growth was larger and we even observed divisions in puzzle-shaped cells (Figure 2F). The area at division in *da1-1eod-2 spch* was approximately threefold larger than in *spch*. Increased cell size and the formation of lobes is typical of epidermal cells that have exited proliferative cell cycles and initiated differentiation [27]. This suggested that cell division arrest may not be a feature associated with differentiation, but rather a change in threshold area. Normal *DA1* and *BB* joint function may either to establish such a threshold area or prolong the duration of cell division across the growing leaf leading to divisions of larger cells.

**Figure 2.**
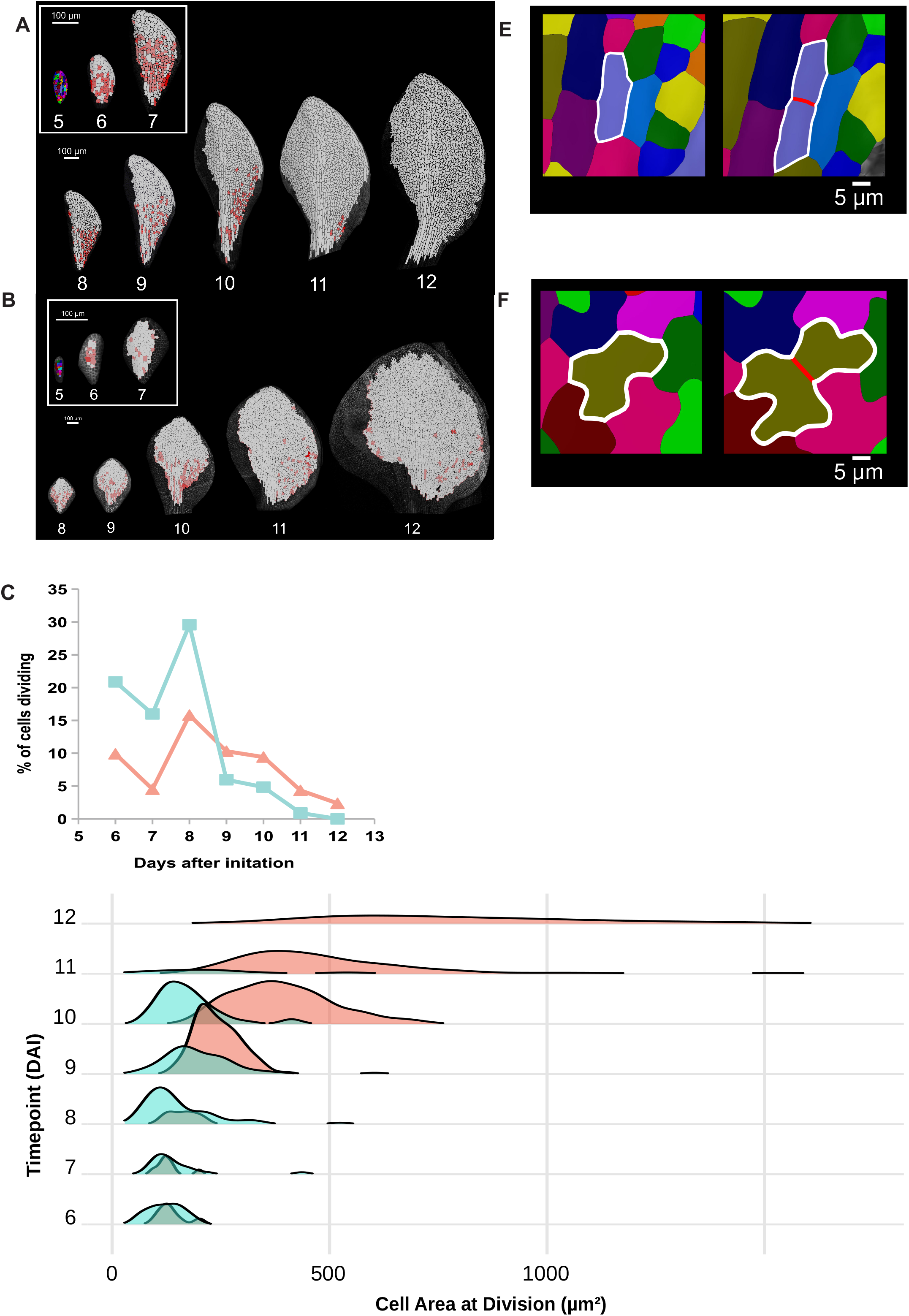
Cell Proliferation and Cell Sizes at Division. A. The image marks cells that divided between the timepoints in red in growing *spch* leaves. B. The image marks cells that divided between the timepoints in red in growing *da1-1 eod1-2 spch* leaves. Note the increased segmented leaf area in this mutant and the continued cell divisions at later timepoints compared to *spch* in panel A. C. The graph shows the percentage of cells divided between each timepoint. Spc is shown as a blue line and *da1-1 eod1-2 spch* as a red line. D. The plot shows the areas of cells at division of spch (blue) and *da1-1 eod1-2 spch* (red). The height of the plots represents the numbers of cells. Note the extended sizes and numbers of dividing cells in *da1-1 eod1-2 spch* compared to *spch* at later stages of leaf growth. E. Image of a typical segmented cell in *spch* (outlined) before and after division. F. Image of a typical segmented cell in *da1-1 eod1-2 spch* (outlined) before and after division. Note the cell lobes compared to *spch* in panel E.

### Divisions occur more distally in *da1-1eod1-2 spch* compared to *spch*

In wild-type leaves, cell proliferation occurs proximally and ceases at the arrest front boundary that moves basipetally as the leaf grows [14,15,27]. To understand how increased cell proliferation seen in *da1-1eod-2 spch* relates to these spatial patterns of cell proliferation, the position of each dividing cell on the leaf was defined according to a single Bezier line along the midrib of the leaf and two sets of normalised positional data for dividing cells at each time point relative to this line were generated (Figure 3A).

**Figure 3.**
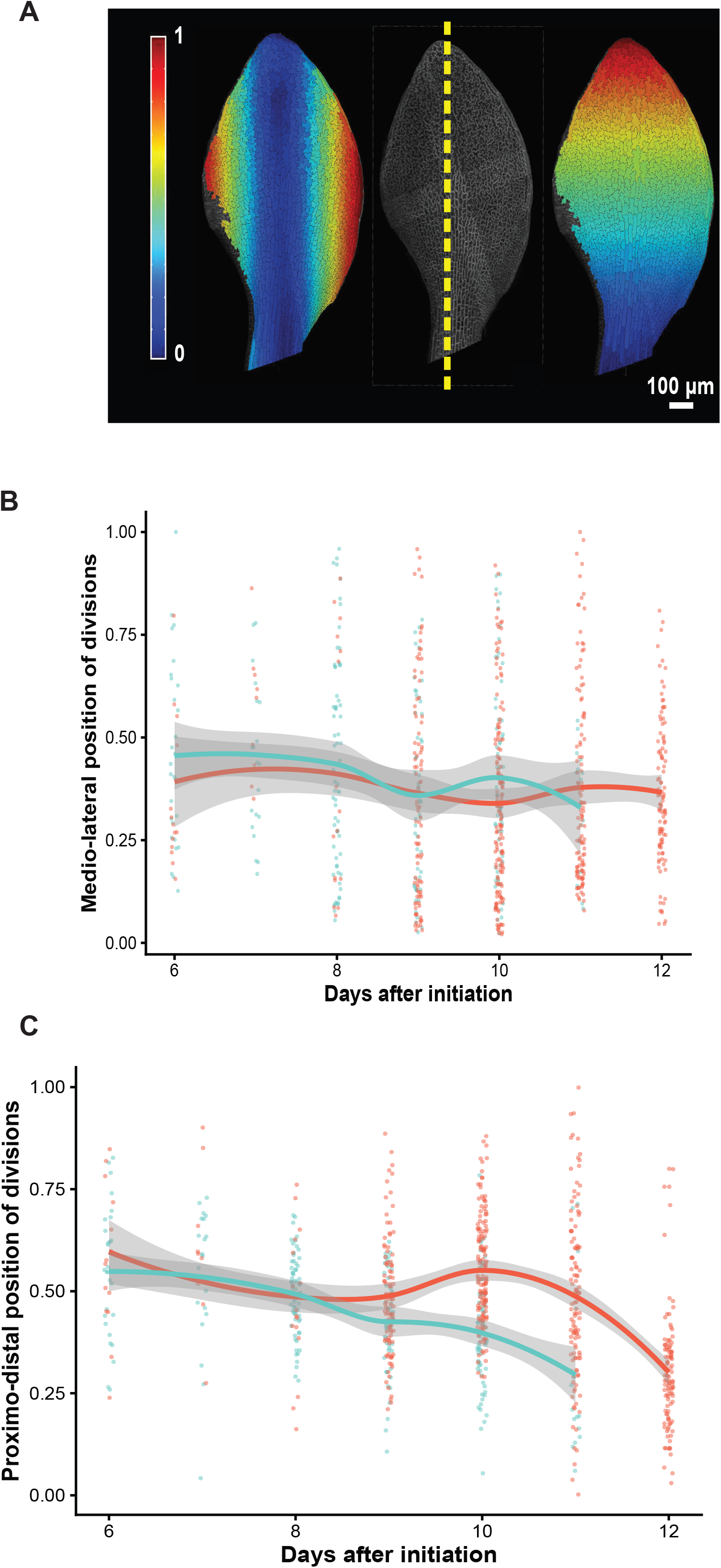
Medio-Lateral and Proximo-Distal Patterns of Cell Division. A. A Bezier line (yellow dashes) demarcates medio-lateral and proximo-distal positions on a segmented leaf. The heatmap shows the axes from 0 (blue) to 1 (red), and the scale bar is 100 μm. B. Medio-lateral patterns of cell division numbers in spch (blue) and *da1-1 eod1-2 spch* (red) lines. Note there is no discernable difference in medio-lateral patterns of division between the mutants. C. Proximo-distal patterns of cell division numbers in spch (blue) and *da1-1 eod1-2 spch* (red) lines. Note the extended pattern of distal cell divisions in *da1-1 eod1-2 spch* compared to *spch*. B&C line of best fit is shown with 95% confidence interval.

Distance from the line gave a medio-lateral position, with 0 being close to the midrib and 1 being close to the margin. Distance along the line gave a proximo-distal distance, with 0 being close to the base and 1 being close to the tip. Using normalised values of each cell division at each time point, relatively even distributions of divisions for both *spch* and *da1-1 eod1-2 spch* in the medio-lateral orientation were observed at all timepoints (Figure 3B). In *spch* between 6 to 8 days, divisions were evenly distributed along the proximo-distal axis, while between 9 to 11 days divisions were generally limited to the proximal region of the leaf (Figure 3B). This was consistent with the appearance of a cell proliferation arrest front appearing to move basipetally during leaf growth [15]. During *da1-1eod1-2 spch* leaf growth, a similar trend of an even distribution of dividing cells along the proximo-distal axis was observed between 6 to 8 DAI, but in contrast, between 9 to 12 DAI significantly more divisions occurred distally compared to *spch* (Figure 3C). Thus, competence to divide persisted into distal regions of *da1-1eod1-2 spch* leaves where cells were larger and more lobed, leading to progressive dissolution of the arrest front.

### Computational modelling of cell division patterns

Our observations from time-lapse analyses from both sets of mutants indicated that cell proliferation occurred in a proximal region of the leaf for a specific time period. In *spch*, cells outside this region do not divide, whereas in *da1-1eod1-2 spch* some cells outside the main proliferative zone keep dividing. Proliferation ceased globally at different times in *spch* and *da1-1eod1-2 spch*.

To explore how the spatial-temporal control of the proliferative zone and proliferation cessation may explain observations, we used a 2D model of dividing cells with uniform, anisotropic growth (descriptive growth). Cell division occurred when cells reached a threshold area, and followed a shortest wall through the centroid rule. An initial template was formed into a leaf shape and populated with cells as described in the Mehtods section[28] (Figure 4A). The amount of growth was set to fit measured leaf lengths. As the leaf grew, cell division occurred when a threshold area was reached. Noise was added to this threshold to approximate the distribution seen in the data. To account for spatial and temporal differences in cell division, two functions were used to control the area threshold for to occur. The first controlled the threshold cell size for divisions as a function of distance from the base of the template. The second was dependent on model time, scaling with the cell size threshold determined by the distance function.

**Figure 4.**
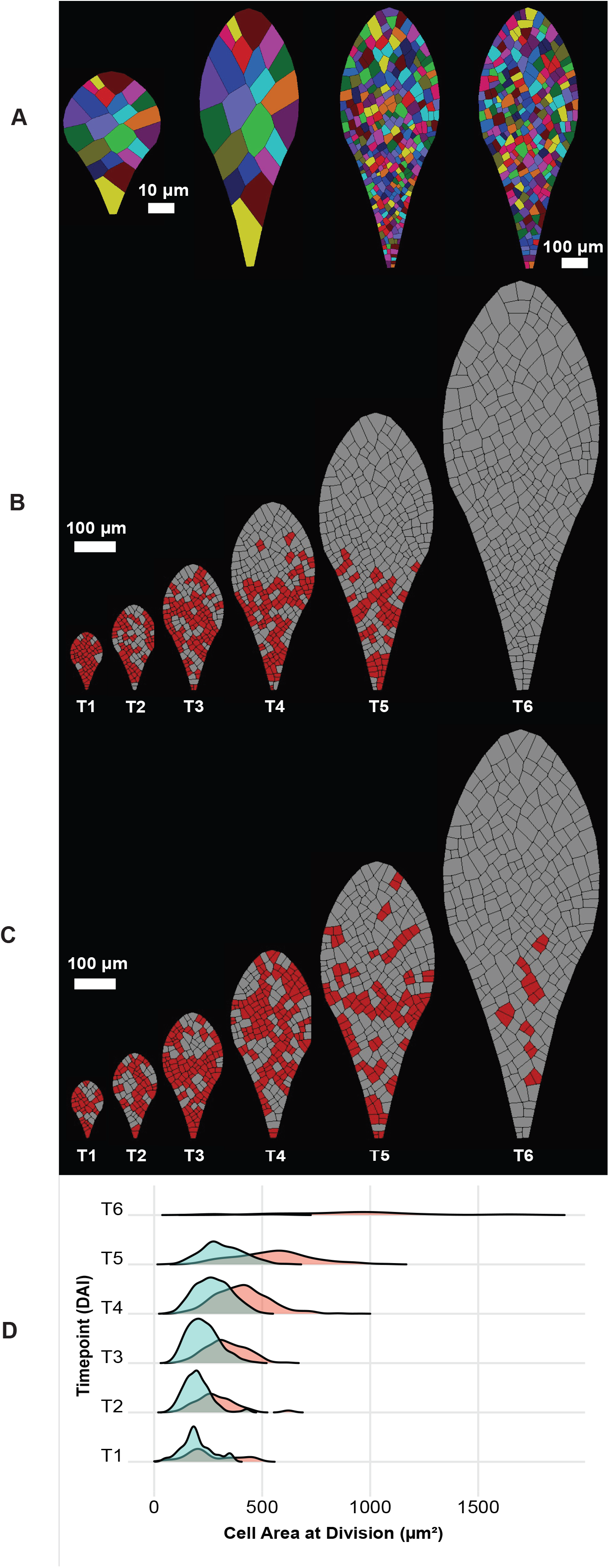
Modelling Spatio-Temporal Patterns of Cell Division. A. Initial template populated with randomly assigned polygons. B. Time series of proliferation patterns with distance and time functions fitting to (B) *spch* and (C) *da1-1 eod1-2 spch* observations. Divisions in between timepoints are highlighted in red. C. Frequency polygon plot showing the areas of cells at division in simulations with distance and time functions fitting to *spch* (blue) and *da1-1 eod1-2 spch* (red). The height of the plots represents the numbers of cells.

Data from growth of *spch* leaves was modelled to establish a baseline. The distance function was initially linear, giving a cell size at division threshold of 200 μm^2^ in a region approximately 200 μm from the base. Beyond 200 μm from the base, there was a steep increase in cell division threshold area that essentially shut down cell division outside the proliferative zone (Supplementary Figure 1A). Since the zone is at a fixed distance from the base, as the leaf grows to 200 μm in length and beyond, it gives the appearance of an arrest front moving basipetally. The time function was also fixed to a scale of 1.0 from the start of the model run, increasing towards the end to model the overall cessation of division as seen in the data at later stages (Supplementary Figure 1A). These model parameters recapitulated the observed dynamics of cell division in *spch* (Supplementary movie 1 and Figure 4B).

Model functions were then changed using experimental observations of *da1-1eod1-2 spch* leaf growth. A much more gradual increase in the threshold size for division in the distance function gave a softer boundary to the proliferative zone and allowed larger, more distal cells to divide (Supplementary Figure 1B). When the time-dependent increase in the division threshold was also made more gradual, proliferation persisted for longer and the overall cessation of cell division was delayed (Supplementary movie 2 and Figure 4C). These models accurately predicted the distributions of cell area at division observed experimentally (compare Figures 4D and 2D). These findings suggest that proliferation arrest is not simply a binary switch. A steep increase in threshold can generate an apparently sharp arrest front, whereas a shallower increase allows occasional divisions in large, lobed cells that appear to show signs of differentiation.

## Conclusions

The growth of determinate organs such as plant leaves is characterised by an initial period of cell proliferation followed by cessation of division, increased cell growth and differentiation. The transition from proliferation to growth is modulated by a wide variety of genes (reviewed by [29] whose activities are thought to function in a spatio-temporal framework [14,15]. Previous work has described the leaf arrest front as a basipetally moving boundary, often viewed as an arrest wave determined by proximo-distal patterning and the shift from proliferation to differentiation [1,10]. Our live imaging and modelling indicated that this front can instead emerge from spatio-temporal control of the cell size threshold for division, rather than from an independently specified boundary. These findings are more consistent with a gradual loss of proliferative competence than with a simple binary switch, because in *spch* a steep increase in threshold generates an apparently sharp front, whereas in *da1-1 eod1-2 spch* large lobed cells can still divide as the threshold rises more gradually. By placing *DA1* and *BB* function in modulating levels of cell proliferation proteins [22,30] upstream of this process, our results reveal how local control of division size can establish organ-scale patterns of proliferation arrest and final leaf size.

## Supporting information

spch growth video

da1-1 eod1-2 spch growth video

## Acknowledgements

We thank Dr Samantha Fox for advice and encouragement. We acknowledge support from the John Innes Foundation Rotation Programme Studentship to NT, the Human Frontiers Science Program grant RPG0067/2021 to RSS, and a Biological and Biotechnological Sciences Research Council Institute Strategic Program grant GEN (BB/P013511/1) to the John Innes Centre (RSS and MWB).

## Key Resources Table

See Supplementary Table 1

## Resource Availability

### Lead Contact

Further information and requests for resources and reagents should be directed to and will be fulfilled by

### Materials Availability

Transgenic lines produced in this study are available upon request.

### Data and Code availability

Confocal images and analyses generated in this paper have been deposited in Zenodo and will be publicly available at the date of publication. DOIs will be in Supplementary Table 1.

Modelling code will be deposited at Zenodo. DOIs will be in Supplementary Table 1. Any additional information required to reanalyse the data reported here is available from the lead contacts.

### Experimental Model and Subject Details

#### Plants and Growth Conditions

All *Arabidopsis thaliana* plants were the Col-0 genotype and are listed in the key resources table. Soil-grown plants were grown in peat-grit mixture in glasshouses with 16h light/8h dark cycle with 120 umol m^2^/sec supplemental fluorescent lighting, 21-23 °C day, 16 °C night temperatures. Plants were grown on Petri dishes on GM agar (0.43% Murashige and Skoog macro- and micro-nutrients and 5mM MES-KOH buffer pH 5.7) in controlled environment growth rooms at 20°C, 16 light/ 8 hr dark cycle.

Transgenic plants were selected by growing T1 plants in peat-grit mix and sprayed with 10 mg/l BASTA each week for 3 weeks. Surviving plants were genotyped using primers listed in the Key Resources Table.

#### Method Details

##### Transgenic Lines

The *pAtML1::mCitrine-RCI2A* plasmid (*pAR169*) was kindly provided by Dr Emily Abrash of Stanford University, with the permission of Dr Adrienne Roeder at Cornell University [31]. The mCitrine fluorescent protein was fused to the plasma membrane localised protein RCI2A and expressed from the epidermal-specific *AtML1* promoter.

##### Plant Transformation, Crossing and Genotyping

*pAtML1::mCitrine-RCI2A* was transformed into Agrobacterium tumefaciens GV3101 by electroporation and selection on LB agar plates with rifampicin (25 ug/ml), gentamycin (25 ug/ml) and spectinomycin (25 ug/ml). Plates were grown at 28°C for 2 days.

*Arabidopsis thaliana* Col-0 plants harbouring the double mutant *da1-1eod1-2* [12] were transformed by floral dip [32] and transgenic lines selected on BASTA as described above. Single copy T-DNA transgenic lines were identified in T1 plants by IDNA Genetics using quantitative PCR of the BASTA gene. These were selfed and T2 plants were assessed for segregation of BASTA resistance (25 ug/ml). A single transgenic line selected for consistent fluorescence (hereafter called *da1-1eod1-2pAR169*) was crossed with the *spch-4* mutant in the Col-0 background (SALK_088595; [25] courtesy of Dr Samantha Fox, John Innes Centre. Lines were genotyped with primers described in Supplementary Table 1 using Edwards buffer purification of crude DNA from seedling leaf tissue ([25,33]. Lines were maintained as *da1-1eod1-2pAR169* homozygotes and *SPC/spch* heterozygotes.[32].

##### Microscopy and Live imaging

Sterilised seeds were sown onto GM agar (0.43% (w/v) Murashige and Skoog, 1% (w/v) sucrose, 0.01% (w/v) inositol, 10ppm (w/v) thiamine, 50ppm (w/v) pyridoxine, 50 ppm (w/v) nicotinic acid, 5mM MES-KOH pH 5.7, 0.9% (w/v) agar). Seeds were stratified in complete darkness for 3 days at 4°C before growth at 20°C, 16 hours light, 8 hours dark. Five days after stratification, seedlings were transferred to sterilised dimpled slides (Sigma-Aldrich. Cat. No. BR475505-50EA) with a droplet of sterilised water and covered by a sterilised coverslip (BDH. Cat. No. 405/0187/35). Seedlings lacking stomata were identified using a Leica SP5 (II) confocal scan head mounted on a Leica DM6000 microscope and transferred to a fresh plate. A sterilised optical live imaging chamber [15,26] was filled with quarter-strength Murashige and Skoog medium (Sigma-Aldrich. Cat. No. M5519) and 0.75% sucrose (Melford. Cat. No. S0809) in a sterile laminar flow hood. Four seedlings with adaxial leaf surfaces positioned uppermost were grown in each chamber and medium constantly circulated. Between imaging time-points, the optical chamber was maintained at 20°C, 16 hours light, 8 hours dark.

Seedlings were imaged every 6 hours for the first 48 hours, every 8 hours for the subsequent 48 hours, every 12 hours for the subsequent 48 hours, and every 24 hours for the final 48 hours. Seedlings were imaged using either a Leica SP5 (II) or a Leica TCS SP8X confocal scan head mounted on a Leica DM6000 microscope, controlled using the Leica LAS AF software. A custom attachment was used to attach the optical imaging chamber to the Leica DM6000 stage. A 20x air objective was used on both microscopes. The argon laser excitation was set to 514 nm, and the fluorescence emission was collected in the 520-600 nm range using a HyD detector.

##### Cellular Growth Analyses

Epidermal cell outlines from successive confocal time points were processed in MorphoGraphX [17,18]. The confocal signal was projected onto a curved surface mesh to generate a 2.5D representation of the leaf epidermis. Cells were segmented on the projected surface using seeded watershed segmentation and segmentation errors were corrected manually where necessary. From the segmented meshes, projected cell area and centroid position were extracted for each cell. Leaf area and cell number were quantified from the segmented epidermis. Areal growth between successive time points was calculated from segmented meshes after lineage assignment.

Cell lineages were assigned between consecutive time points by parent labeling in MorphoGraphX. This enabled time-lapse analysis of cell proliferation and growth in developing leaves, as in previous live-imaging studies [15,34]. Parent cells at time t were matched to corresponding cells at time t + Δt, and where division had occurred, both daughter cells were assigned the same parent label. Division events were therefore identified when one parent cell at time t corresponded to two daughter cells at the following time point. The proportion of proliferating cells was calculated as the number of cells that had divided between two time points expressed relative to the total number of segmented cells at the later time point. Cell area at division was estimated as the sum of the projected areas of the two daughter cells immediately after division.

For positional analysis of proliferating cells, a single Bézier line was drawn along the midrib of each leaf. The position of each dividing cell was then expressed using two normalized coordinates relative to this line: proximo-distal position, defined by distance along the line from leaf base to tip, and medio-lateral position, defined by perpendicular distance from the line to the leaf margin. These coordinates were used to quantify the spatial distribution of divisions across the growing lamina.

##### Leaf Growth Modelling

Simulation models of a growing leaf with cell division were created using the MorphoDynamX framework (www.MorphoDynamX.org) and the vlab modeling environment (https://www.algorithmicbotany.org/virtual_laboratory)[28] . Cells in the models were represented as polygonal faces defined by edges and vertices. Positions were stored in vertices, with the origin located at the bottom edge of the leaf in the center. Growth was anisotropic, with more growth along the proximo-distal axis of the leaf in a ratio of 5:4. To implement growth, vertex positions were scaled uniformly at each time step. Model time was divided into 6 sections of equal size of 0.39 units, where the length of the template at each point was similar to the leaf length found in the data (T1 = 0.95, T2 = 1.34, T3 = 1.73, T4 = 2.12, T5 = 2.51, T6 = 2.9). The models started with a reference shape that approximated the early leaf shape, which as in the experiments, was partially occluded by the cotyledons. This shape was then subdivided into cells by using a shortest wall algorithm. Polygons were divided such that the new edge was as short as possible while passing through the centroid of the cell. Cells were divided until all were below an initial threshold area of 150 μm2.

Two functions were used to control the threshold for the division area. These spline functions can be edited interactively with the vlab funcedit program, or by manually changing control point positions in the files themselves. The first function controlled the threshold cell size for divisions as a function of distance from the base of the template. Since vlab functions are typically defined such that values are between 0 and 1 in both axes, scaling factors were used in the model to scale the input and output values. In the threshold function based on distance, the input value was scaled by 0.001 so that 1.0 in the x-axis of the function approximates the size of the leaf before division arrest, and the output scaled by 2000 so that 0.1 represents an initial starting threshold area of 200 μm2. The second function controls threshold area based on global model time, and acts as a multiplier for the first. Its input value is scaled by 0.35 so that 1.0 represents the complete run time of 2.9. The output was scaled by 10.0, so that the initial value of 0.1 represents no scaling (i.e. a threshold of 200.0 μm2).

Significant noise is always observed in the cell size at division. In order to simulate noise, it is not sufficient to simply add noise to the division threshold, as cells are tested against it at each time step causing noise to be cancelled out, especially when the time step is small. To mitigate this we introduced a specific noise value for each daughter cell at the time of division. With each cell a multiplier against the division threshold was stored and given a Gaussian distribution about 1.0 with the sigma specified as a model parameter (0.3). When a daughter cell grew, this multiplier was then applied against the threshold determined by the functions to control division size.

To visualize proliferation, a time was associated with daughter cells at division that was set to the current global model time. Cells which had a time within a window approximating the experimental time points were considered to be recently divided and visualized in red.

##### Quantification and Statistical Analyses

Plots were generated in R using the packages ggplot2. Frequency plots in figure 2D were generated using the package ggridges with smoothed ridgelines representing data density. Comparisons between genotypes at each timepoint was performed using the Mann-Whitney Test, showing no significant difference at DAI 6-8, and significant differences at DAI 9-11 (DAI 6 *p* = 0.357, DAI 7 *p* = 1, DAI 8 *p* = 0.082, DAI 9 *p* < 0.001, DAI 10 *p* < 0.001, DAI 11 *p* < 0.001). To determine the magnitude of overlap between two groups, Cliff’s Delta test was used at each timepoint (a value of 0 is a perfect overlap, a value of 1 is no overlap and a value >0.474 is a significant difference in overlap). We find the data highly overlaps at DAI 6-8 (DAI 6 *d* = 0.179, DAI 7 *d* = 0.000, DAI 8 *d* = 0.296) and has a low overlap at DAI 9-11 (DAI 9 *d* = 0.427, DAI 10 *d* = 0.912, DAI 11 *d* = 0.723).

Frequency plots in 4G were generated using the same method. Comparisons between model timepoints using the Mann-Whitney Test showed no significant difference (T1 *p* < 0.001, T2 *p* < 0.001, T3 *p* < 0.001, T4 *p* < 0.001, T5 *p* < 0.001, T6 *p* < 0.001). The magnitude of overlap between timepoints using Cliff’s Delta test showed overlaps that broadly matched the data (T1 *d* = 0.236, T2 *d* = 0.736, T3 *d* = 0.592, T4 *d* = 0.596, T5 *d* = 0.647, T6 *d* = 0.875).

## Supplementary Material

Supplementary Figure 1

A. Distance (left panel) and time (right panel parameters for *spch* growth models

B. Distance (left panel) and time (right panel parameters for *da1-1 eod1-2 spch* growth models

C. Cell areas at division generated for *spch* and (D) *da1-1 eod1-2 spch* by the growth models

Supplementary Table 1. Key Resources Table

Supplementary Movies

