## Supplementary Table 1 for "The leaf cell arrest front is established by spatio-temporal control of cell sizes at division"

**Supplementary Table 1. Key Resources Table**

| REAGENT or RESOURCE | SOURCE | IDENTIFIER |
| --- | --- | --- |
| Antibodies | | |
| Bacterial and virus strains | | |
| Agrobacterium tumefaciens GV3101 | widely available | Holsters et al 1980 |
| Escherichia coli DH5-α | ThermoFisher | Cat. 18265017 |
| Biological samples |  |  |
| Chemicals, peptides, and recombinant proteins | | |
| BASTA (d-l glufosinate) | Sigma-Aldrich | Cat. 45520 |
| Murashige and Skoog complete medium | Sigma-Aldrich | Cat. M5534 |
| Sucrose | Melford | Cat. 50809 |
| Silwet L-77 | Fisher | Cat. NC0138454 |
| MES Buffer | Sigma-Aldrich | Cat. M0164 |
| Rifampicin | Sigma-Aldrich | Cat. R3501 |
| Gentamycin | Sigma-Aldrich | Cat. G1264 |
| Spectinomycin | Sigma-Aldrich | Cat. 54014 |
| Critical commercial assays | | |
| iDNA Genetics copy number RT PCR Service | <https://www.norwichresearchpark.com/partner/idna-genetics> |  |
| SYBR Green PCR master mix | ThermoFisher | Cat.4367659 |
| Deposited data | | |
| Confocal images and code will be available from Zenodo |  |  |
| Experimental models: Cell lines | | |
| Experimental models: Organisms/strains | | |
| Arabidopsis thaliana Col-0 | George Redei | ABRC CS60000 |
| Arabidopsis thaliana Col-0 da1-1eod1-2 | Li et al 2008 | N/A |
| Col-0 *pAtML1::mCitrine-RCI2A* | Roeder A et al 2010 | N/A |
| Col-0 *da1-1bb pAtML1::mCitrine-RCI2A* | This study | N/A |
| Col-0 *spch-4* | MacAlister et al 2007 | N/A |
| Oligonucleotides | | |
| GACACCATGCAATGCCAACC. Li et al 2008 | Sigma-Aldrich | da1-1 F |
| CTTTGAGCCTCATCCACGCA. Li et al 2008 | Sigma-Aldrich | da1-1 R |
| ATTTTGCCGATTTCGGAAC T-DNA left border | Sigma-Aldrich | LB1.3 |
| GAGCGATGCATCTCTAACCAC. SALK_04519 LP | Sigma-Aldrich | eod1-2 L |
| AGTAGGAACAGAAAGCAGGGG. SALK_045169 RP | Sigma-Aldrich | eod1-2 R |
| TATGAGGGACTCGCATTCATC SALK_078595 LP | Sigma-Aldrich | spch L |
| AAAACAAATTCGTTTGCTCCC. SALK_078595 RP | Sigma-Aldrich | spch R |
| Recombinant DNA | | |
| Plasmid *pAtML1::mCitrine-RCI2A* | Roeder A et al 2010 | E. Abrash, Stanford |
| Software and algorithms | | |
| Leica Application Suite X (LAS X) | Leica |  |
| FIJI |  |  |
| MorphoGraphX (MGX V) |  |  |
| R |  |  |
| Customised R scripts will be availble from Zenodo |  |  |
