## Supplementary figures and images for "The leaf cell arrest front is established by spatio-temporal control of cell sizes at division"

### Supplementary Figure 1

**Supplementary Figure 1**

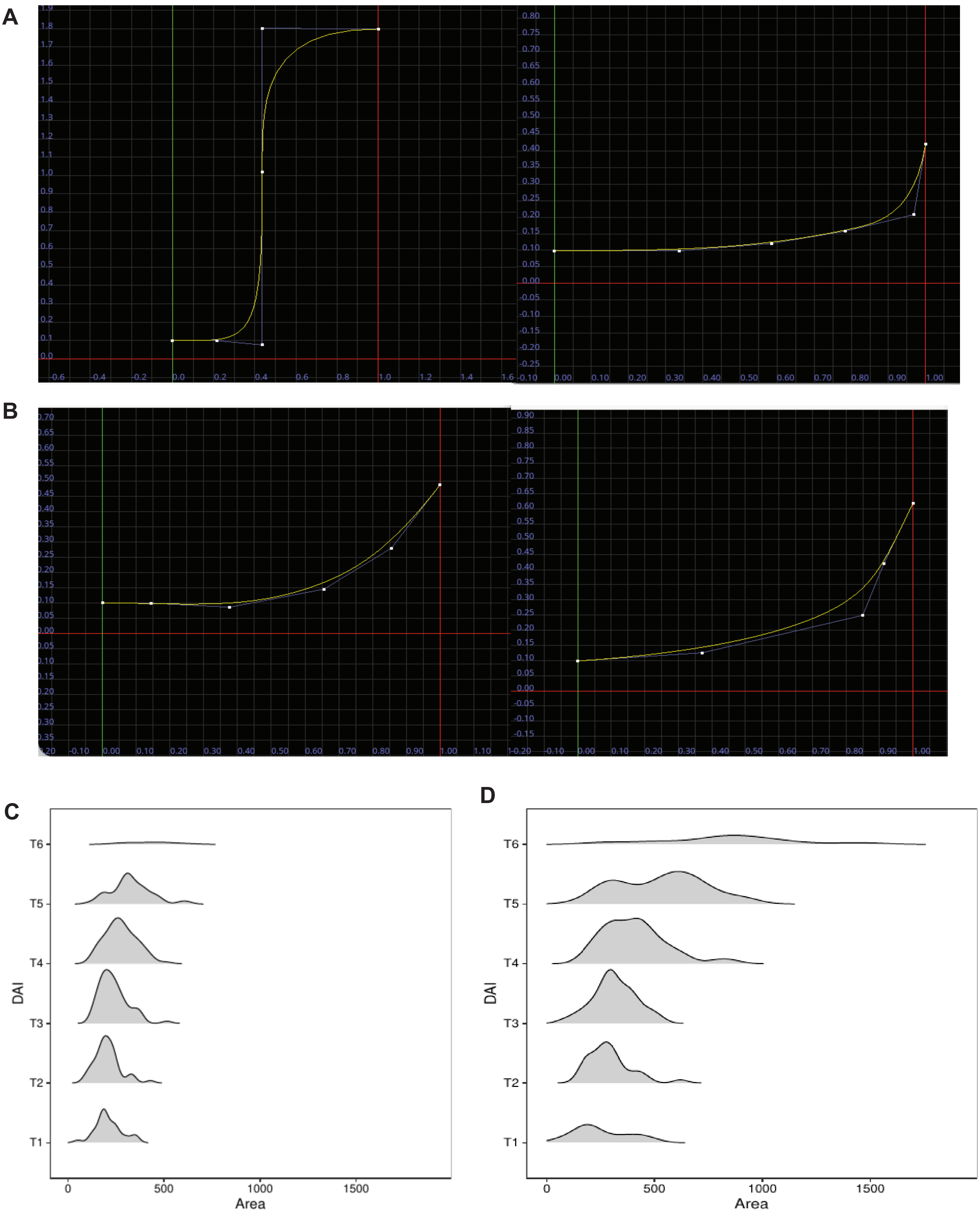
